# BaSiCPy: Scalable and Robust Shading Correction for Optical Microscopy Images

**DOI:** 10.64898/2026.04.28.721386

**Authors:** Yu Liu, Yohsuke T. Fukai, Santiago Cano-Muniz, Victor Perez, Mihail Ivilinov Todorov, Germán Camargo Ortega, Timothy Morello, Alexander Jovanovic, Dirk Loeffler, Johannes C. Paetzold, Xun Xu, Nan Ma, Lorenz Lamm, Timm Schroeder, Lucas Boeck, Denis Schapiro, Nicholas Schaub, Ali Ertürk, Jianxu Chen, Carsten Marr, Tingying Peng

**Affiliations:** School of Computation, Information and Technology, Technical University of Munich, Munich, Germany; Nonequilibrium Physics of Living Matter Laboratory, RIKEN Pioneering Research Institute, 2-2-3 Minatojima-minamimachi, Chuo-ku, Kobe, Hyogo 650-0047, Japan; Nonequilibrium Physics of Living Matter RIKEN Hakubi Research Team, RIKEN Center for Biosystems Dynamics Research, 2-2-3 Minatojima-minamimachi, Chuo-ku, Kobe, Hyogo 650-0047, Japan; Department of Biomedicine, University of Basel, Basel, Switzerland; Institute for Computational Biomedicine, Faculty of Medicine, Heidelberg University and Heidelberg University Hospital, Heidelberg, Germany; Translational Spatial Profiling Center (TSPC), Heidelberg, Germany; GSK, Cellzome GmbH, Heidelberg, Germany; Institute for Intelligent Biotechnologies (iBIO), Helmholtz Center Munich, Neuherberg, Germany; Institute for Stroke and Dementia Research (ISD), University Hospital, LMU Munich, Munich, Germany; Department of Biosystems Science and Engineering, ETH Zurich, 4058 Basel, Switzerland; Department of Physiology & Pharmacology, SUNY Downstate Health Sciences University, Brooklyn, New York, USA; Institute of Active Polymers and Berlin-Brandenburg Center for Regenerative Therapies, Helmholtz-Zentrum Hereon, Teltow, Germany; Institute of Chemistry and Biochemistry, Freie Universität Berlin, Berlin, Germany; Helmholtz AI, Helmholtz Munich, Neuherberg, Germany; Institute of Pathology, Faculty of Medicine, Heidelberg University and Heidelberg University Hospital, Heidelberg, Germany; IFOM ETS - the AIRC Institute of Molecular Oncology, Milan, Italy; National Center for Advancing Translational Sciences, National Institutes of Health, Bethesda, MD, USA; Center for Advancing Medical AI, Novagen Research Fund (NGRF), North Bethesda, MD, USA; Axle Research and Technology, North Bethesda, MD, USA; Graduate School of Systemic Neuroscience (GSN), University Hospital, LMU Munich, Munich, Germany; Munich Cluster for Systems Neurology (SyNergy), Munich, Germany; Deep Piction, Munich, Germany; School of Medicine, Koç University, İstanbul, Turkey; Leibniz-Institut für Analytische Wissenschaften – ISAS – e.V., Dortmund, Germany; Computational Health Center & Helmholtz AI, Helmholtz Munich, Neuherberg, Germany; Department of Medicine III, LMU University Hospital, LMU Medizin, LMU Munich, Germany; Department of Physics, Ludwig-Maximilians-Universität München, Munich, Germany; German Cancer Consortium (DKTK), partner site Munich, Germany; Munich Center for Machine Learning (MCML), Munich, Germany

## Abstract

Quantitative fluorescence microscopy is frequently confounded by spatially varying illumination and temporal intensity drift. Although BaSiC is a widely adopted retrospective correction method, it can fail when foreground content is strongly correlated across images—a common regime in time-lapse, tiled and volumetric acquisitions—and its application often requires manual parameter tuning that limits reproducibility and scalability. We introduce BaSiCPy, a foreground-aware implementation of BaSiC that improves illumination profile estimation under correlated foreground structures, provides automatic hyperparameter selection and accelerates large-scale processing through GPU support. BaSiCPy is distributed as an open-source Python package with graphical and programmatic interfaces, facilitating integration into contemporary bioimage analysis workflows.

## Main

Modern microscopy increasingly produces large, heterogeneous datasets, including tiled multiplexed tissue images, volumetric light-sheet acquisitions, and long time-lapse recordings, in which quantitative analyses depend on stable intensity calibration across space and time. Yet spatially varying illumination, detector inhomogeneity and temporal baseline drift introduce systematic biases that appear as stitching seams in mosaics^1^ and spurious trends in longitudinal measurements^2^. Illumination correction is therefore essential for reliable quantification in high-throughput and large-scale imaging studies.

To this end, retrospective approaches estimate illumination directly from the acquired images and are widely used when prospective calibration images are unavailable or impractical^2–4^. Among them, BaSiC models an image sequence as a smoothly varying flatfield (and optional darkfield) together with a temporally varying baseline, enabling joint correction of shading and drift^1,2,5–7^. However, BaSiC, together with other retrospective methods that rely on separating illumination from sample signal, can produce biased illumination estimates when foreground structures remain highly correlated across frames or slices, as in slowly varying time-lapse recordings or volumetric stacks with continuous anatomy. In such settings, persistent biological structure can leak into the estimated illumination profile, leading to over-correction and distortion of quantitative readouts.

We present BaSiCPy, an open-source Python implementation that extends BaSiC for robust use in contemporary microscopy workflows. BaSiCPy mitigates biases caused by correlated foreground structures through segmentation mask–guided reweighting during illumination estimation, and it removes dataset-specific manual tuning through automatic hyperparameter selection. BaSiCPy further supports scalable processing through GPU acceleration and provides accessible interfaces, including a Napari plugin (**Extended Data Fig. 1**) and guided notebook workflows, for integration into modern bioimage analysis pipelines.

Following BaSiC, BaSiCPy models a shading-corrupted image sequence *I*^′^(*t, x*) as (**Fig. 1A**):

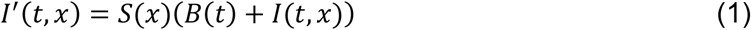

where *S*(*x*) represents the spatially inhomogeneous flatfield, *B*(*t*) is the spatially constant yet temporally varying baseline, and *I*(*t, x*) is the underlying biological signal, respectively. Because this factorization is underdetermined and cannot be solved directly, BaSiCPy assumes that the flatfield *S*(*x*) is shared across the sequence and varies smoothly in space, while the biological signal *I*(*t, x*) is comparatively more structured and less smooth. These priors enable accurate estimation of both spatial and temporal illumination profiles^2,3^. As a result, BaSiCPy effectively corrects illumination heterogeneity across space, reducing uneven shading and stitching seams in multi-tile mosaics (**Fig. 1B–C**), and across time by stabilizing baseline intensity in long-term acquisitions (**Fig. 1D**). Moreover, BaSiCPy, compared to BaSiC, incorporates foreground-aware illumination estimation and automatic hyperparameter selection, improving robustness under challenging acquisition conditions.

**Fig. 1.**
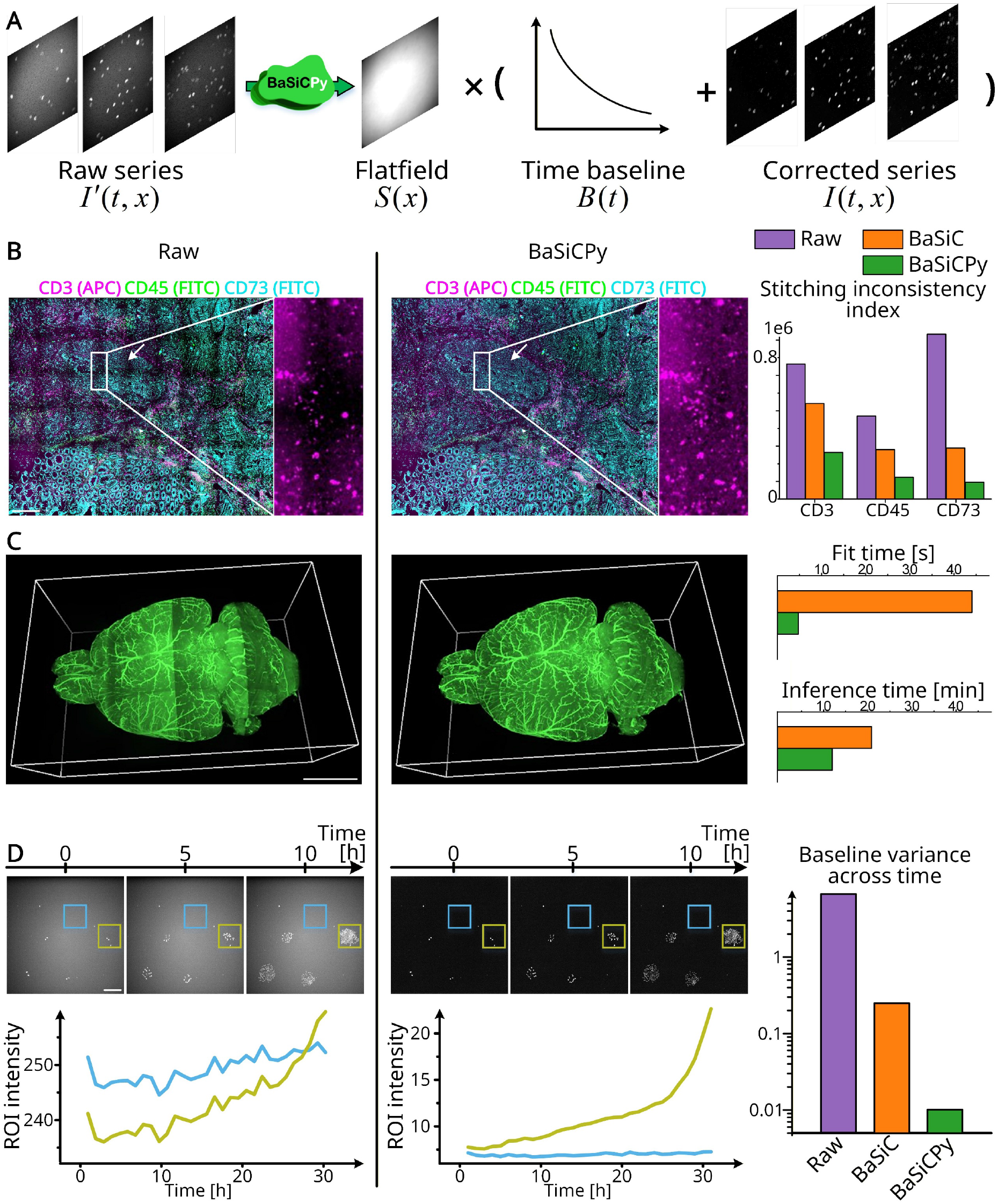
BaSiCPy enables robust flatfield estimation and quantitative stabilization across imaging modalities. (A) Conceptual model of BaSiCPy. A shading-corrupted image sequence *I*′(*t, x*) is decomposed into a spatial flatfield *S*(*x*) and a temporally varying baseline *B*(*t*), allowing recovery of the corrected signal *I*(*t, x*). (B–D) Representative correction results across three imaging modalities. In each case, raw data (left) are compared to BaSiCPy-corrected results (middle), followed by quantitative evaluation (right). (B) Multiplexed imaging. Raw mosaics of a colorectal cancer tissue exhibit tile-border intensity discontinuities due to uneven illumination. BaSiCPy reduces stitching artifacts, as quantified by a stitching inconsistency index, primarily due to improved flatfield estimation enabled by Autotune-selected hyperparameters. (C) 3D tiled volumetric imaging. Uneven illumination across tiles of a large-scale whole-mouse-brain vasculature dataset (raw dataset size ≈500 GB) is visible in raw projections and largely removed after correction. Leveraging GPU acceleration, BaSiCPy substantially outperforms the original BaSiC implementation: ∼9 times faster in flatfield fitting, and ∼2× faster in correcting the entire volume. All runtime benchmarks were conducted on a standard laptop equipped with an Intel Core Ultra 9 185H processor (16 cores and 22 logical processors), 32 GB of RAM, and an NVIDIA GeForce RTX 4060 Laptop GPU with 8 GB of memory. (D) Time-lapse imaging with correlated foreground. Raw data of cytokine-supported murine hematopoietic stem and progenitor cells exhibit temporal baseline drift and background intensity fluctuations. BaSiCPy stabilizes background intensity while preserving biological signal dynamics. Background baseline variance across time is substantially reduced compared to raw data and BaSiC through foreground-aware reweighting that mitigates correlated foreground bias.

For example, in cyclic multiplexed imaging of a colorectal cancer tissue section acquired as multi-channel mosaics (6×9 tiles per channel) using the MACSima platform (**Fig. 1B**), BaSiCPy markedly reduced stitching-induced grid artifacts compared to BaSiC. Over time, BaSiCPy also stabilizes baseline intensity in long-term widefield time-lapse imaging of live hematopoietic progenitor cells (**Fig. 1D**), where gradual signal drift and photobleaching typically introduce substantial temporal variability.

To address the correction bias due to correlated foreground, where consistently elevated intensities at fixed spatial locations are misinterpreted as local illumination inhomogeneity (dotted circles, **Fig. 2A**), BaSiCPy incorporates a foreground-aware reweighting strategy that uses optional segmentation masks to down-weight foreground pixels during illumination estimation (**Fig. 2A**). To quantitatively validate the effect of foreground awareness, we simulated a shading-free cytoplasm time-lapse sequence with limited cell mobility and high foreground occupancy (up to 83.7%), introducing strong spatial and temporal correlation between foreground and illumination (**Methods**). Spatially uneven illumination was imposed using a prospectively acquired dye-based flatfield together with exponential baseline decay to model photobleaching. Both BaSiC and mask-free BaSiCPy exhibited severe flatfield errors (Γ^S > 300%; **Extended Data Fig. 2**). In contrast, foreground-aware BaSiCPy accurately recovered the ground-truth flatfield (Γ^S < 5%) and stably corrected the temporal baseline drift. Comparable flatfield accuracy was obtained using either ground-truth or threshold-based masks, demonstrating robustness to imperfect foreground segmentation (**Extended Data Fig. 2**).

We next evaluated BaSiCPy on a multi-channel time-lapse widefield fluorescence microscopy dataset of primary blood progenitors labeled with distinct surface markers (**Extended Data Fig. 3**). Across markers with distinct expression patterns and foreground densities, conventional correction exhibited channel-dependent behaviors, including structured flatfield artifacts or incomplete baseline stabilization. In contrast, foreground-aware BaSiCPy consistently recovered smooth illumination profiles and yielded temporally stable background intensities across all tested channels. We further evaluated BaSiCPy on a whole-mouse-brain vasculature dataset acquired using UltraMicroscope II (**Fig. 1C, Extended Data Fig. 4**). In this image, consecutive sampling of limited axial slices introduced strong inter-slice correlation due to continuous tissue structures. While conventional correction by BaSiC required larger sampling windows to achieve stable illumination estimation (**Extended Data Fig. 4A**), foreground-aware BaSiCPy maintained consistent flatfield recovery even when using substantially fewer consecutive slices (**Extended Data Fig. 4B**). As a result, BaSiCPy yielded lower stitching inconsistency indices at all tested sampling numbers, significantly better than BaSiC (Wilcoxon signed-rank test, p<0.05).

Beyond correlated foreground bias, a second practical challenge in illumination correction is achieving robust performance across diverse imaging modalities, sample types, and acquisition conditions. In practice, parameters that control the smoothness of the estimated illumination profile often require dataset-specific tuning to balance overfitting and under-correction. Within the BaSiC framework, this sensitivity is primarily governed by the flatfield smoothness regularization term (smoothness_flatfield, *λ*_*s*_), which controls the trade-off between suppressing foreground leakage and preserving genuine illumination gradients. As illustrated in Fig. 2B (bottom), small *λ*_*s*_ values yield unstable flatfield estimates contaminated by high-frequency structure from foreground signals, whereas excessively large *λ*_*s*_ values produce overly smooth profiles that fail to capture true illumination inhomogeneity. The optimal values of *λ*_*s*_ vary substantially across datasets (**Extended Data Table 1**), making it difficult to predetermine a single suitable value of *λ*_*s*_.

**Fig. 2.**
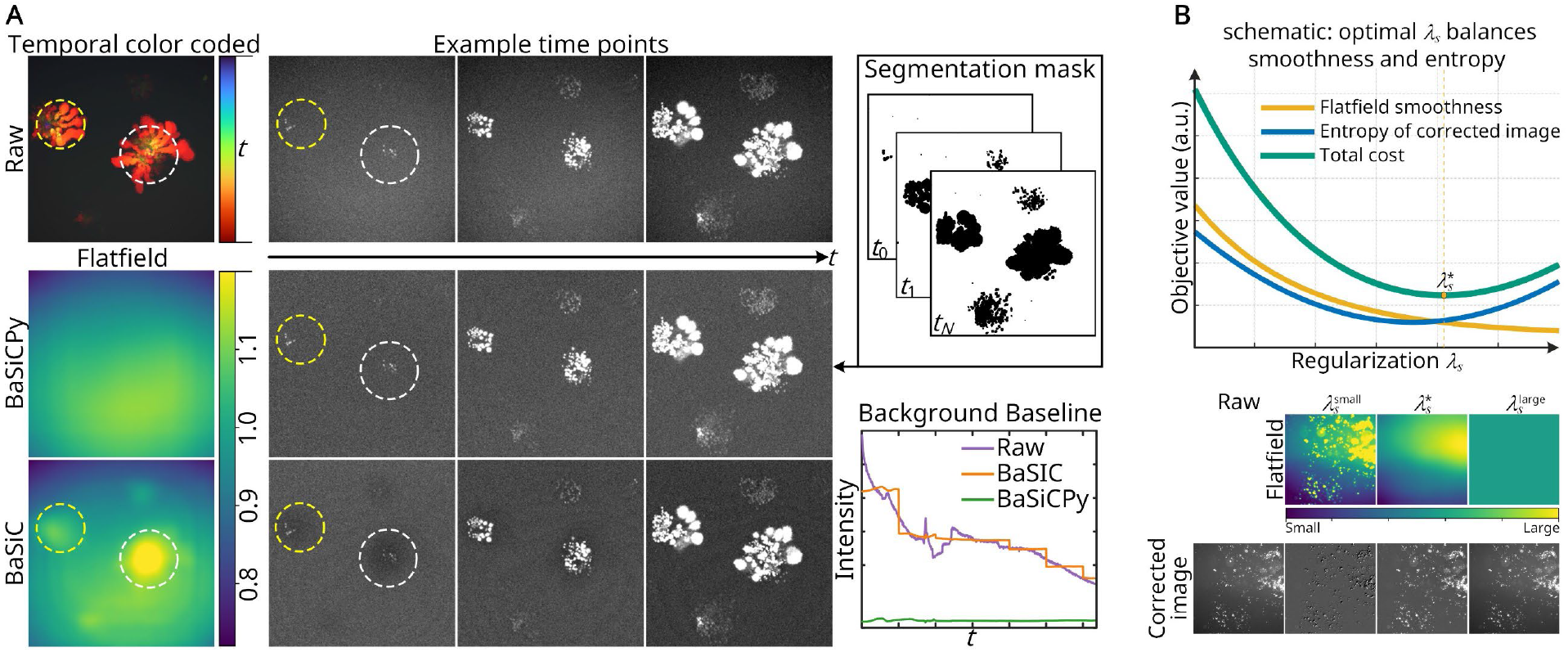
BaSiCPy mitigates correlated foreground bias and enables automatic hyperparameter selection. (A) Foreground-aware correction. In time-lapse imaging with limited cell motility, conventional methods, such as BaSiC, misinterpret static foreground as illumination (dotted circles). BaSiCPy uses a segmentation mask to down-weight foreground pixels during flatfield estimation, recovering a clean illumination profile. The resulting baseline stabilization (bottom) outperforms raw data and BaSiC. (B) Automatic hyperparameter tuning (Autotune). The flatfield smoothness parameter *λ*_*s*_ controls the trade-off between over-smoothing (high *λ*_*s*_) and foreground leakage (low *λ*_*s*_). Autotune selects the optimal 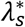 minimizing a cost function that balances flatfield smoothness and the entropy of the corrected image (top).

Conceptually, the optimal *λ*_*s*_ balances flatfield smoothness against correction quality, which motivates the Autotune strategy in BaSiCPy for automatic parameter selection. Specifically, Autotune selects 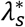 by minimizing a cost function that combines a Fourier-based flatfield smoothness term and an entropy-based correction quality term (**Fig. 2B**, top). The optimum is identified using a deterministic two-stage grid search.

To evaluate this behavior in a realistic high-content imaging scenario, we applied Autotune to an antimicrobial single-cell testing (ASCT) time-lapse dataset^8^ acquired across multiple wells and imaging positions (**Extended Data Fig. 5**), where propidium iodide (PI) fluorescence was tracked over time under different drug conditions, resulting in heterogeneous intensity distributions across wells and frames. Sweeping *λ*_*s*_ produced substantially different flatfield estimates for each well, reflecting variations in foreground density, background levels, and illumination patterns. In contrast, the Autotune-selected 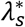 consistently yielded well-adapted flatfield profiles without manual adjustment. Importantly, the corrected time-lapse sequences exhibited markedly stabilized baseline intensities across wells, whereas the raw signals showed pronounced temporal decay and variability. This fully automatic setting is particularly advantageous for high-throughput screening workflows, where large numbers of wells and conditions make manual tuning of illumination parameters impractical. Notably, all results reported in this study were obtained using Autotune-selected parameters without manual hyperparameter tuning.

To further evaluate BaSiCPy’s utility for downstream quantification, we used the ASCT dataset to compare the killing efficiency of three chemically distinct antibiotics—azithromycin (AZM), clofazimine (CFZ), and cefoxitin (FOX)—a central goal in high-content antimicrobial screening. As shown in **Fig. 3** (left, “Without BaSiCPy”), raw data do not permit such a comparison due to drug-dependent artifacts: spatial illumination non-uniformity (halo effect, well-border shadowing), drug autofluorescence (CFZ overlaps PI emission at 618 nm), and temporal variability (photobleaching, lamp fluctuations, temperature drift). Each antibiotic suffers from a different mixture of these biases, making cross-comparison invalid—e.g., the normalized live fraction (relative to time 0) erroneously exceeds 1.0 (reaching 1.2), which is biologically impossible. After BaSiCPy correction (**Fig. 3**, right, “With BaSiCPy”), artifacts are substantially reduced: the flatfield corrects spatial gradients, and the temporal baseline compensates for drift. As quantified in the bottom panel of **Fig. 3**, all three antibiotics now yield normalized live fractions within the biologically plausible range of 0–1, following the expected killing dynamics. BaSiCPy thus enables reliable cross-antibiotic comparison within the same experiment.

**Fig. 3.**
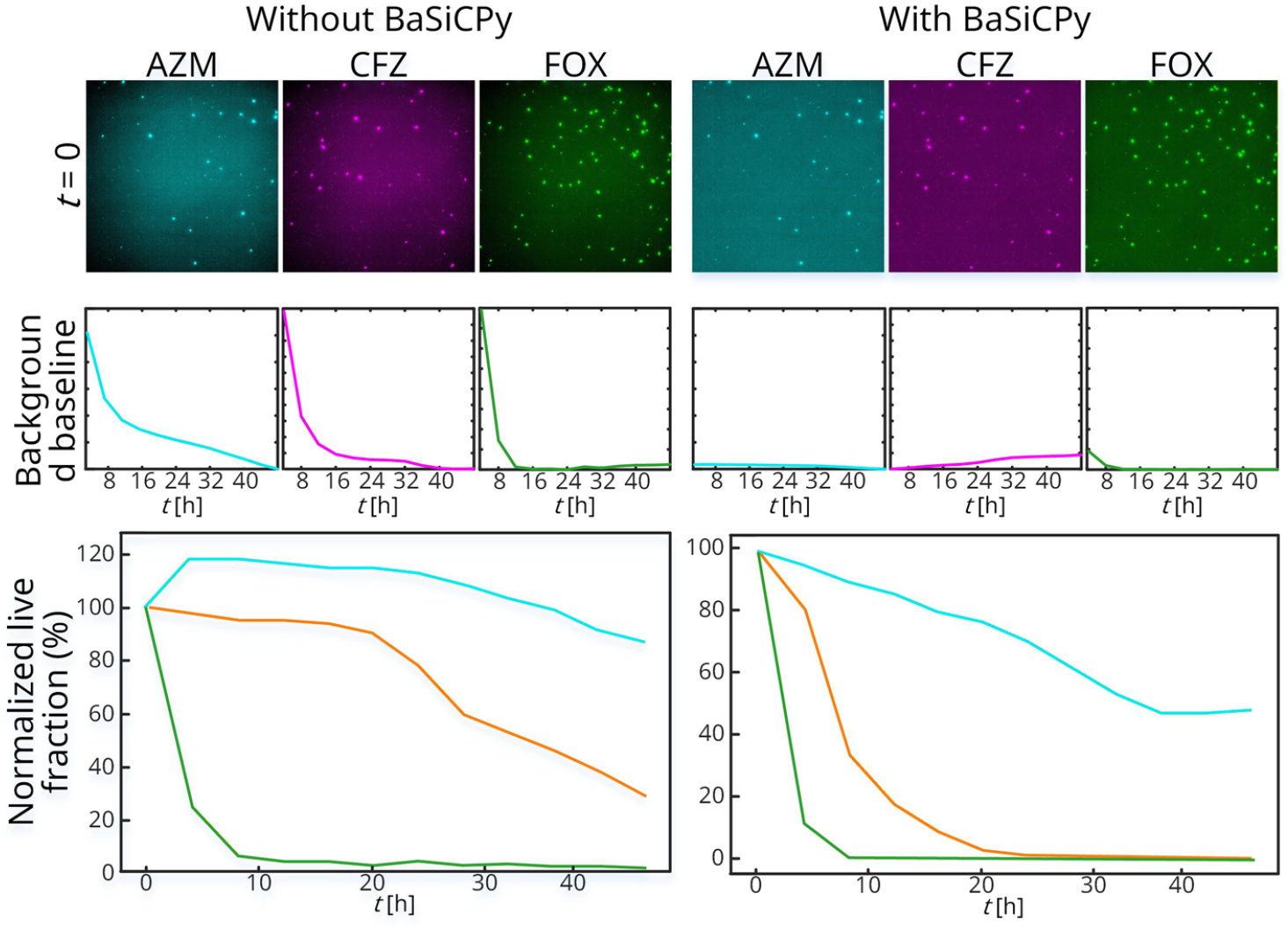
BaSiCPy’s application to antimicrobial single-cell testing (ASCT). Raw data of Mycobacterium abscessus treated with azithromycin (AZM), clofazimine (CFZ), and cefoxitin (FOX) exhibit drug-dependent illumination issues, making cross-comparison invalid. After BaSiCPy correction, all three antibiotics yield normalized live fractions within [0, 1] and follow the expected killing dynamics, enabling reliable cross-antibiotic comparison.

Beyond algorithmic improvements, BaSiCPy clarifies several previously ambiguous design choices in BaSiC to improve practical usability across diverse imaging modalities. Specifically, components such as darkfield estimation and intensity sorting are retained as optional features rather than default steps. Specifically, in modern microscopy workflows, additive darkfield contributions are often negligible or already absorbed into residual backgrounds, making explicit darkfield estimation unnecessary in most cases. Nevertheless, rare scenarios such as physical contamination or fixed-pattern defects on the sensor (e.g., dust, stains, or dead pixels) may still benefit from darkfield correction, and BaSiCPy therefore preserves this functionality as an advanced option (**Extended Data Fig. 6**,**Extended Data Fig. 7**). Similarly, while intensity sorting can stabilize optimization in static datasets, it may distort temporal fidelity in time-lapse imaging. BaSiCPy consequently disables sorting by default and recommends its use only in modality-specific contexts. In addition, BaSiCPy provides modality-aware usage clarifications for complex imaging paradigms. For example, in cyclic multiplexed imaging workflows where marker and autofluorescence channels are acquired separately, the background signal is spatially heterogeneous and does not conform to a quasi-homogeneous darkfield model. In such cases, illumination correction should be applied to the raw marker and autofluorescence channels independently rather than to their difference image (**Extended Data Fig. 8**). More detailed modality-specific recommendations are maintained in the online documentation [https://basicpy.readthedocs.io/en/latest/] and will be continuously updated to reflect emerging community practices and diverse microscopy workflows.

In summary, BaSiCPy extends the BaSiC framework into a robust, open-source tool for illumination correction in contemporary microscopy. By introducing segmentation mask-guided reweighting, it overcomes the long-standing failure of retrospective methods when foreground structures remain correlated across frames or slices, as in time-lapse recordings and volumetric stacks of continuous anatomy. The Autotune strategy removes the need for dataset-specific manual tuning of the flatfield smoothness parameter, enabling consistent performance across diverse modalities, sample types, and acquisition conditions. Combined with GPU acceleration and accessible napari and notebook interfaces, these advances make reliable illumination correction practical at the scale and heterogeneity of modern high-throughput and large-scale imaging studies. As demonstrated across multiplexed tissue mosaics, multi-channel and volumetric acquisitions, and high-content antimicrobial screening, BaSiCPy enables quantitative readouts that would otherwise be confounded by spatial, temporal, and foreground-dependent biases, supporting trustworthy downstream analysis in modern bioimage workflows.

## Acknowledgments

D.S. is supported by the German Federal Ministry of Education and Research (BMBF 01ZZ2004); the Ministry for Science, Research and Science Baden-Württemberg “AI Health Innovation Cluster” and “MULTI-SPACE”; research funding from Cellzome, a GSK company, the Bruno and Helene Jöster Stiftung, IFOM ETS Institutional Funds and the Multi-dimensionAI project (CZS-Project number: P2022-08-101) was made possible by funding from the Carl-Zeiss-Stiftung. This work is supported by the Health + Life Science Alliance Heidelberg Mannheim and received state funds approved by the State Parliament of Baden-Württemberg. We thank Dr. Nadya Panagides and Dr. Maximilian Wirth for providing the multiplexed imaging dataset used in this study. C. M received funding from the European Research Council under the European Union’s Horizon 2020 Research and Innovation Programme (grant agreements 866411, 101113551, and 101213822) and support from the Hightech Agenda Bayern. This work was supported through the Bavarian AI Foundation Model Initiative, funded by the Bavarian State Ministry of Science and the Arts (StMWK).

## Declaration of interests

Denis Schapiro (D.S.) reports funding from GSK and received fees/honoraria from Alpenglow, ariadne.ai, GSK, Immunai, Lunaphore, and Noetik.

## Data availability

Datasets used to evaluate BaSiCPy are publicly available on the Zenodo repository (https://doi.org/10.5281/zenodo.6334810). The repository includes representative cell culture data, multi-channel time-lapse fluorescence microscopy datasets, brightfield time-lapse recordings, and whole-slide brain imaging data. The remaining samples are either publicly available at the link provided or can be requested from the original investigators

## Code availability

BaSiCPy is a pip-installable Python package and is available at the following GitHub repository: https://github.com/peng-lab/BaSiCPy, with documentation at https://basicpy.readthedocs.io/en/latest/. Tutorials regarding the use of BaSiCPy as Notebooks and Napari plugins can be found at https://basicpy.readthedocs.io/en/latest/tutorials.html.

## Methods

### Recap of BaSiC

Before introducing BaSiCPy, we start from a simplified form of the low-rank–sparse optimization underlying BaSiC for estimating the spatial flatfield *S*(*x*) and the per-frame baseline *B*(*t*) in Eq. (1) from an observed image sequence *I*^′^(*t, x*) *∈* ℝ^*T*×*M*×*N*^, which is either a time-lapse dataset with *T* frames of 2D slices of size *M* × *N*, or a 3D volumetric dataset with *T* axial planes. Specifically, BaSiC formulates illumination estimation as the following optimization problem:

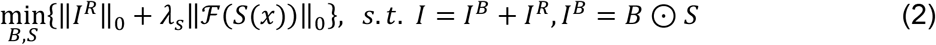

Where *⊙* is the outer product, *I ∈* ℝ^*T*×(*M*×*N*)^ is the measurement matrix obtained by vectorizing each slice in *I*^′^(*t, x*) as a row, *S ∈* ℝ^1×(*M*×*N*)^ is the vectorized flatfield, *B ∈* ℝ^*T*×1^ is the per-frame baseline and their outer product *I*^*B*^ is a rank 1 matrix, *I*^*R*^ *∈* ℝ^*T*×(*M*×*N*)^ is the residual matrix, *λ*_*s*_ is the balancing parameter and is set to be:

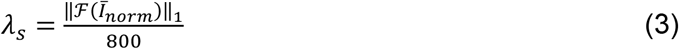

where 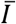 is the mean of *I* cross the *T* dimension and later normalized, 800 is empirically set. In Eq. (2), two priors are used: (i) spatial smoothness of the flatfield *S*(*x*), as smooth functions are typically sparse in the Fourier transformed domain, i.e., ‖ℱ (*S*(*x*))‖_0_, and (ii) the residual *I*^*R*^ is sparse relative to the low-rank part as it collects mainly the foreground contributions (i.e., specimen signal is uncorrelated along *T* axis). Since the optimisation of a L0-norm is NP-hard, Eq. (2) is replaced via:

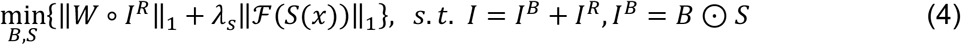

where the sparsity of ℱ (*S*(*x*)) is imposed via standard L1 norm because normal L1-norm is good enough when the matrix has a small number of non-zero coefficients (high sparsity) and ℱ (*S*(*x*)) indeed is highly sparse as a flatfield is usually smooth with most of its energy concentrated in low-frequency bands. In contrast, the sparsity of *I*^*R*^ is less predictable, as it largely depends on the image content, and thereby its sparsity is imposed via an iterative reweighting L1 norm and *W ∈* ℝ^*T*×(*M*×*N*)^ is the reweighting matrix, *∘* is an element-wise product. *W* are initially set to be 1 and iteratively updated by:

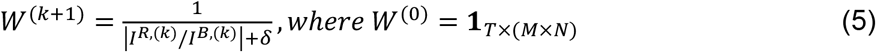

where *δ* > 0 is introduced to provide stability. In practice, BaSiC estimates the flatfield and baseline through an iterative reweighted optimization procedure based on Eq. (4). The detailed optimization scheme is described in the original BaSiC publication^2^ and summarized in **Extended Data Fig. 9**.

### BaSiCPy: foreground-aware uneven illumination correction

A key limitation of existing illumination correction approaches, including BaSiC, is inaccurate flatfield estimation when foregrounds are highly correlated. In such cases, *I*^*R*^ is no longer sparse but may instead exhibit low-rank structure aligned with *I*^*B*^. The reweighting *W ∝ I*^*B*,(*k*)^/*I*^*R*,(*k*)^ further over-weights these low-rank structures, causing the foreground signal to be absorbed into *S* and biasing the estimation of *S* and *B* in the end.

To mitigate this bias, BaSiCPy integrates an optional segmentation mask Ω *∈* {0,1}^*T*×(*M*×*N*)^ (1 for background, 0 for foreground) into the reweighting scheme of Eq. (4). We keep the BaSiC objective and solver but only modify how the reweighting matrix *W* in the iterative reweighting L1 norm is initialized and updated. Specifically, instead of initializing *W* directly with Ω, BaSiCPy starts from a unit matrix, and applies down-weighting to pixels masked by Ω for numerical stability:

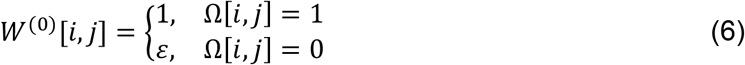

where *ε*> 0 and by default *ε* = 0.1. Later at the *k*^*th*^ iteration, BaSiCPy updates *W* by first reusing Eq. (5) and then down-weighting entities masked by Ω:

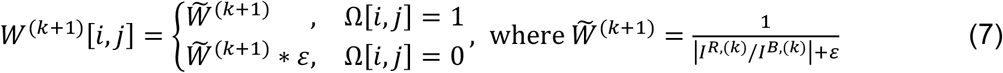

All remaining steps (updates of *S, B*, multiplier *Y*, and penalty *µ*) follow the BaSiC algorithm. Iteratively, correlated foreground signal remains in *I*^*R*^, as the corresponding thresholds are nearly negligible when updating *I*^*R*,(*k*+1)^ to boost sparsity (see line 17 in **Extended Data Fig. 9**). The analyses that use this feature are listed in **Extended Data Table 2**.

### BaSiCPy: automatic hyperparameter tuning

In practice, we observed that the balancing parameter *λ*_*s*_ has the largest impact on the correction outcome. Specifically, it controls the smoothness of the estimated flatfield *S*(*x*) by penalizing its sparsity in the Fourier domain. If *λ*_*s*_ is set too low, the estimated flatfield becomes unstable and may retain high-frequency structures originating from foreground objects, leading to overfitting. In contrast, if *λ*_*s*_ is set too high, the flatfield will be overly smoothed, failing to capture the illumination inhomogeneities. Thus, an appropriate *λ*_*s*_ is crucial for achieving reliable correction performance.

Unfortunately, the optimal *λ*_*s*_ is highly data-dependent. It varies with the degree of uneven illumination, the strength of foreground structures, and so on. These factors differ substantially across experimental setups, imaging modalities, and even acquisition sessions of the same microscope. As a result, manual tuning of *λ*_*s*_ is not only labor-intensive but also difficult to generalize, motivating the need for an automatic hyperparameter, i.e., *λ*_*s*_, selection strategy.

Hence, we propose an Autotune strategy to automatically determine *λ*_*s*_, which first considers the smoothness of the estimated flatfield *S*(*x*). Specifically, we measure how “non-smooth” *S*(*x*) is, by counting the number of significant high-frequency coefficients in its 2D discrete cosine transform (DCT2):

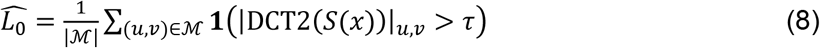

where mask 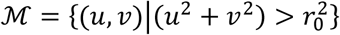 is set in the DCT domain to select high-frequency coefficients, |ℳ| is the number of coefficients in ℳ, *1*(·) is the indicator function that equals 1 if the coefficient amplitude at position (*u, v*) exceeds amplitude threshold *τ*. Here, the amplitude threshold *τ* and the radius parameter *e*_0_ are empirically set to 0.1 and 10, respectively. As a result, a smaller value of 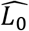 indicates a smoother flatfield *S*(*x*) with fewer local oscillations, whereas larger values suggest that foreground texture or noise has leaked into *S*(*x*). To avoid over-penalizing inevitable small high-frequency DCT coefficients introduced by noise, we adopt a hinge-style penalty:

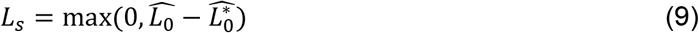

which tolerates a baseline level 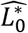 (10^−3^ by default). However, *L*_*s*_ alone tends to favor overly smooth flatfield *S*(*x*), potentially compromising the ability to correct illumination inhomogeneity. To counteract this risk, we introduce an additional entropy-based term. In practice, we observe that shading effects broaden the histogram of the measured image sequence, leading to a more dispersed distribution, whereas a well-corrected volume exhibits a more compact histogram with concentrated background intensities. This observation can be quantified by computing the Shannon entropy of the histogram:

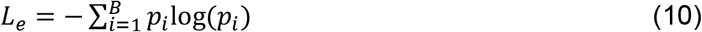

where 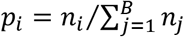 is the normalized probability of the *i*-th histogram bin, and *n*_*i*_ is the number of pixels falling into the *i*-th histogram bin. Here, the histogram is computed within an intensity interval of *v*_*min*_, *v*_*min*_ + *v*_*range*_. To ensure fair comparison across different candidate values of *λ*_*s*_, the histogram range *v*_*range*_ must be fixed: a narrower range would artificially lower the entropy by concentrating the distribution, while a wider range would increase it by dispersing the counts. Therefore, we first fix *v*_*range*_ = *v*_*max*_− *v*_*min*_ × *v*_min _*f*actor_ × *v*_*range _factor*_ from an initial coarse correction by setting *λ*_*s*_ based on Eq. (3), where *v*_*min*_ and *v*_*max*_ are taken as the 1% and 99% quantiles of the corrected image, respectively. In practice, we empirically set *v*_min _*f*actor_ = 0.6 and *v*_range _*f*actor_ = 1.5, which is a balance between excluding extreme outliers and retaining the main intensity distribution. During Autotune, the lower bound *v*_*min*_ is updated as the 0.01-quantile of each candidate correction and scaled by *v*_min _*f*actor_ to ensure that the histogram always aligns with the current background distribution.

Overall, the Autotune strategy minimizes the following cost function, a balance between the Fourier-based smoothness term and the entropy-based compactness term:

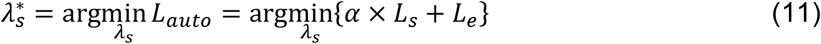

where *α* is a scaling coefficient empirically set to 10^4^. This large value is chosen to match the magnitude of *L*_*s*_ (typically 10^−3^ − 10^−2^) with that of *L*_*e*_ (typically on the order of 1 − 10). Specifically, since BaSiCPy is relatively insensitive to moderate changes of *λ*_*s*_, a simple deterministic two-stage grid search is sufficient to find a near-optimal 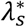 with low overhead and full reproducibility. We first evaluate candidates on a fixed coarse log-grid {0.01, 0.1,0.5,2,8,10}. The coarse argmin is then bracketed with the better neighbor (unique interior neighbor if at boundaries) and refined on a fine log-grid drawn from {0.01, 0.1,0.5,0.75,1,1.25,1.5, 1.75,2,2.5,3,4,5,6,7,8,10}, restricted to the bracket. The searching range can also be adapted by users. This procedure is deterministic and scales linearly with the number of grid evaluations. A detailed summary of the Autotune strategy is provided in **Extended Data Fig. 10**.

### Simulating the effect of uneven illumination and time-lapse baseline drift

We first generated a shading-free and drift-free time-lapse image sequence using an established fluorescence simulation tool^9^. To mimic cell division, the simulation was initialized with one single cell in the first frame, and one additional cell was introduced in each subsequent frame. To reproduce the low cell motility of many stem cell types (e.g., embryonic or hematopoietic stem cells), we enforced strong frame-to-frame correlation by assigning a high probability of cell clustering in the simulation, thereby preventing newly generated cells from migrating away from their birthplace. This resulted in a sequence of 200 2D images (256 × 256 pixels) with fluorescently labeled cytoplasm.

Next, to introduce spatially uneven illumination, we applied a predefined multiplicative flatfield to the shading-free series. To make our simulation as realistic as possible, we used a flatfield obtained prospectively from a concentrated dye solution. In addition, we simulated temporal baseline drift by imposing an exponentially decaying background intensity to mimic photobleaching. The resulting simulated image sequence is shown in **Extended Data Fig. 2**.

### Application scope and best practices in multiplexed cyclic imaging

Multiplexed cyclic imaging generates multi-tile, multi-channel datasets acquired across multiple staining and imaging cycles^10–12^. This acquisition paradigm gives rise to a characteristic data structure that aligns naturally with the modeling assumptions and computational design of BaSiCPy, making it well-suited for flatfield estimation in this modality.

First, multiplexed datasets typically comprise tens to hundreds of spatially distributed tiles per channel, which promotes effective suppression of correlated foreground structures and facilitates robust flatfield recovery under the low-rank plus sparse decomposition model adopted by BaSiCPy. Second, the moderate tile count per channel (typically 50–100 tiles) provides sufficient sampling for accurate flatfield estimation while maintaining favorable convergence properties. Third, as channels and cycles are independent, flatfield estimation and correction can be parallelized across channels and/or cycles, enabling scalable deployment on large, multiplexed imaging studies.

As a best practice, we do not recommend using BaSiCPy for direct estimation of autofluorescence or darkfield baselines in multiplexed datasets. The algorithm is designed to correct for an additive background baseline, i.e. a quasi-homogenous background across space. Yet autofluorescence in multiplexed imaging is not a background “baseline” since each tile’s background varies in both intensity and in space. Hence, the heterogeneous backgrounds and foregrounds across tiles make BaSiCPy unsuitable for calculating the darkfield of the whole acquisition. Fortunately, many of the multiplexed imaging platforms also acquire reference images of the autofluorescence^11,12^. Thus, a more appropriate approach is to apply flatfield correction to both marker signal and autofluorescence images followed by their weighted arithmetic subtraction; this procedure yields optimal results (**Extended Data Fig. 8**).

#### Canonical multiplexed imaging processing pipeline

For multiplexed cyclic imaging datasets, each channel is processed using the following canonical pipeline:

1. For each marker channel, flatfields are estimated from the raw marker tiles using BaSiCPy.
2. For the corresponding autofluorescence reference channel, flatfields are independently estimated using BaSiCPy.
3. All tiles are flatfield corrected using their corresponding flatfields according to:

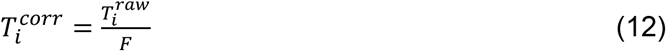

where 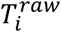 and 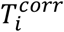 denote the raw and corrected tiles, and *F* denotes the estimated flatfield for the corresponding channel.
4. After flatfield correction, tiles from all channels and cycles are registered and stitched using the ASHLAR algorithm^13^, with DAPI channels serving as spatial references.
5. For each stitched channel, autofluorescence is removed by:

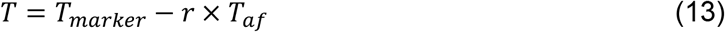

where *r* denotes the ratio of exposure times between marker and autofluorescence acquisitions.

### Quantification of stitching-induced grid artifacts

To quantify residual tile-to-tile intensity inconsistencies after illumination correction, multi-tile mosaics were reconstructed using the Fiji Grid/Collection stitching plugin with identical registration parameters. To isolate intensity mismatch effects, stitching was performed using three different overlap merging rules: minimum intensity (*I*_*min*_), maximum intensity (*I*_*max*_), and linear blending (*I*_*lin*_).

Residual grid artifacts were quantified as:

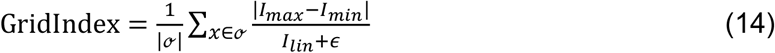

evaluated within tile-overlap regions.

When illumination correction is insufficient, tile borders exhibit intensity discontinuities, resulting in increased divergence between minimum and maximum overlap merging. Lower index values indicate improved inter-tile intensity consistency and reduced stitching artifacts.

### Quantification of similarity between the estimated and the ground-truth flatfields

We quantify the error between the estimated flatfield *S*^*est*^ and the ground-truth *S*^*gt*^ as Γ^*S*^, which is defined as:

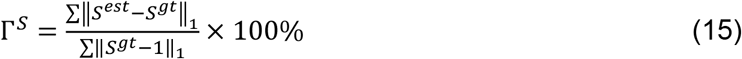

where ‖·‖_1_ denotes the L1 norm, and the constant 1 represents a uniform flatfield. A lower value of Γ^*S*^ indicates higher similarity between the estimated flatfield and the ground-truth.

### Background baseline computation

To quantitatively evaluate the temporal baseline drift in both simulated and real datasets, we computed the background baseline intensity for each time-lapse sequence. For a given image stack *I ∈* ℝ^*T*×*M*×*N*^ with a total of *T* frames of size *M* × *N*, and a binary segmentation mask Ω *∈* {0,1}^*T*×*M*×*N*^(1 for background, 0 for foreground, see previous section for details), the background baseline *b ∈* ℝ^*T*^ was defined as the spatial mean intensity over background pixels:

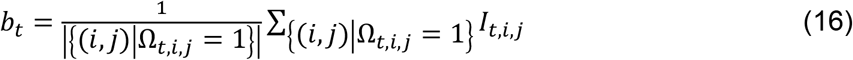

where *b*_*t*_ denotes the mean background intensity of the *t*-th frame.

### Data acquisition

#### Multiplexed imaging

Multiplexed imaging data were acquired using the MACSima platform (Miltenyi Biotec). A colorectal cancer tissue section (4 μm thick) was imaged in cyclic multiplexed mode. For each channel, a mosaic of 6×9 tiles (54 tiles in total) was acquired to cover the region of interest. Dedicated autofluorescence reference images were acquired in each cycle using the same optical configuration as the corresponding marker channels. Exposure times for marker and autofluorescence acquisitions were recorded and used for weighted autofluorescence subtraction during post-processing.

#### Antimicrobial single-cell testing (ASCT) time-lapse dataset

Time-lapse microscopy data were obtained from antimicrobial single-cell testing (ASCT) experiments of Mycobacterium abscessus (strain ATCC 19977)^8^. Bacteria were cultured in Middlebrook 7H9 broth supplemented with 10% OADC and 0.05% Tween-80 to mid-log phase, washed, and embedded in 0.8% low-melting-point agarose containing propidium iodide (PI, 2 μg mL^−1^) for viability assessment.

For demonstration in this study, three representative antibiotics were selected from the ASCT experiments: azithromycin (AZM), cefoxitin (FOX), and clofazimine (CFZ), each applied at 20× minimum inhibitory concentration (MIC). While the full ASCT workflow typically involves a larger panel of compounds and conditions, these drugs were chosen to illustrate illumination variability.

Live-cell imaging was performed for 48 h with image acquisition every 4 h (12 time points) using a Nikon Ti2 inverted microscope equipped with a Prime 95B camera and a SpectraIII/Celesta multi-laser illumination system. Images were acquired at 40× magnification with a pixel size of 0.183 μm, yielding a field of view of 294 μm × 294 μm and image dimensions of 1608 × 1608 pixels (16-bit). Both brightfield (x40BF) and fluorescence (Cy3/PI; excitation 537 nm, emission 618 nm) channels were recorded at multiple positions per well.

Detailed procedures for ASCT image acquisition and downstream analysis, including background correction, bacterial segmentation, viability classification, single-cell tracking, and quantification of time–kill kinetics, have been described previously in Jovanovic et al^8^.

#### Hematopoietic progenitor live-cell multicolor time-lapse imaging

Time-lapse microscopy data were acquired from cytokine-supported murine hematopoietic stem and progenitors cultured in phenol-red- and riboflavin-free IMDM (Seraglob, Cat# M2217) supplemented with 20% BIT serum substitute (STEMCELL Technologies, Cat# 09500), 2 mM GlutaMAX (Thermo Fisher Scientific, Cat# 35050-038), 50 U ml^−1^ penicillin, 50 μg ml^−1^ streptomycin (Thermo Fisher Scientific, Cat# 31350010), 50 μM 2-mercaptoethanol (Thermo Fisher Scientific, Cat# 31350010), 100 ng ml^−1^ murine SCF, 100 ng ml^−1^ murine THPO, with or without 10 ng ml^−1^ murine IL3. Cells were maintained at 37 °C under 5% O_2_ and 5% CO_2_.

For lineage and state detection, the medium was supplemented with fluorochrome-conjugated antibodies, including Ly-6A/E (Sca-1; BioLegend, Cat# 108102, RRID: AB_313339, used at 50 ng/ml), which was labeled in-house with ATTO514 and used at 50 ng ml^−1^, Ly-6A/E (Sca-1-Alexa Fluor 488; eBioscience, Cat# 108116), FcyR-BV480 (BD OptiBuild™ BV480 Rat Anti-Mouse CD16/CD32; Cat# 746324), CD41-PE (CD41a Monoclonal Antibody (eBioMWReg30 (MWReg30)), PE, eBioscience™; Cat# 12-0411-81) and LysoBriteNIR (AAT Bioquest; Cat# 22641), CD16/32 BV421 (BioLegend, Cat# 101332, RRID: AB_2650889, used at 5 ng/ml), CD71 BV480 (BD Biosciences, Cat# 746468, RRID: AB_2743770, used at 20 ng/ml), CD48 Alexa Fluor 594 (BioLegend, Cat# 103436 (discontinued), RRID: AB_2617019, used at 50 ng/ml), CD150 Alexa Fluor 647 (BioLegend, Cat# 115918, RRID: AB_2239178, used at 50 ng/ml). Cells were immobilized on u-Slide VI 0.4 channel slides (IBIDI) coated with 5 μg ml^−1^ anti-CD43 IgM monoclonal antibody (ThermoFisher Scientific, Cat# 14-0431-82, RRDI: AB_467241) in PBS for 1 hour at room temperature.

##### Ethical statement

All experiments were done according to Swiss federal law and institutional guidelines of ETH Zurich, approved by the local animal ethics committee Basel-Stadt (approval number 2655).

##### Research Animals

Experiments were conducted with 12–16-week-old, male C57BL/6J mice from Janvier Labs purchased from The Jackson Laboratory and acclimatized for at least 1 week before experiments. Mice were housed in improved hygienic conditions in individually ventilated cages (2-5 mice per cage max.) with environmental enrichment, and an inverse 12 h day– night cycle in a temperature (21 ± 2 °C) and humidity (55 ± 10%) controlled room with ad libitum access to standard diet and drinking water at all times. The well-being of all mice was monitored regularly by animal facility caretakers, and mice were euthanized in case of symptoms of pain and/or distress.

## Extended Data Figures

**Extended Data Fig. 1.**
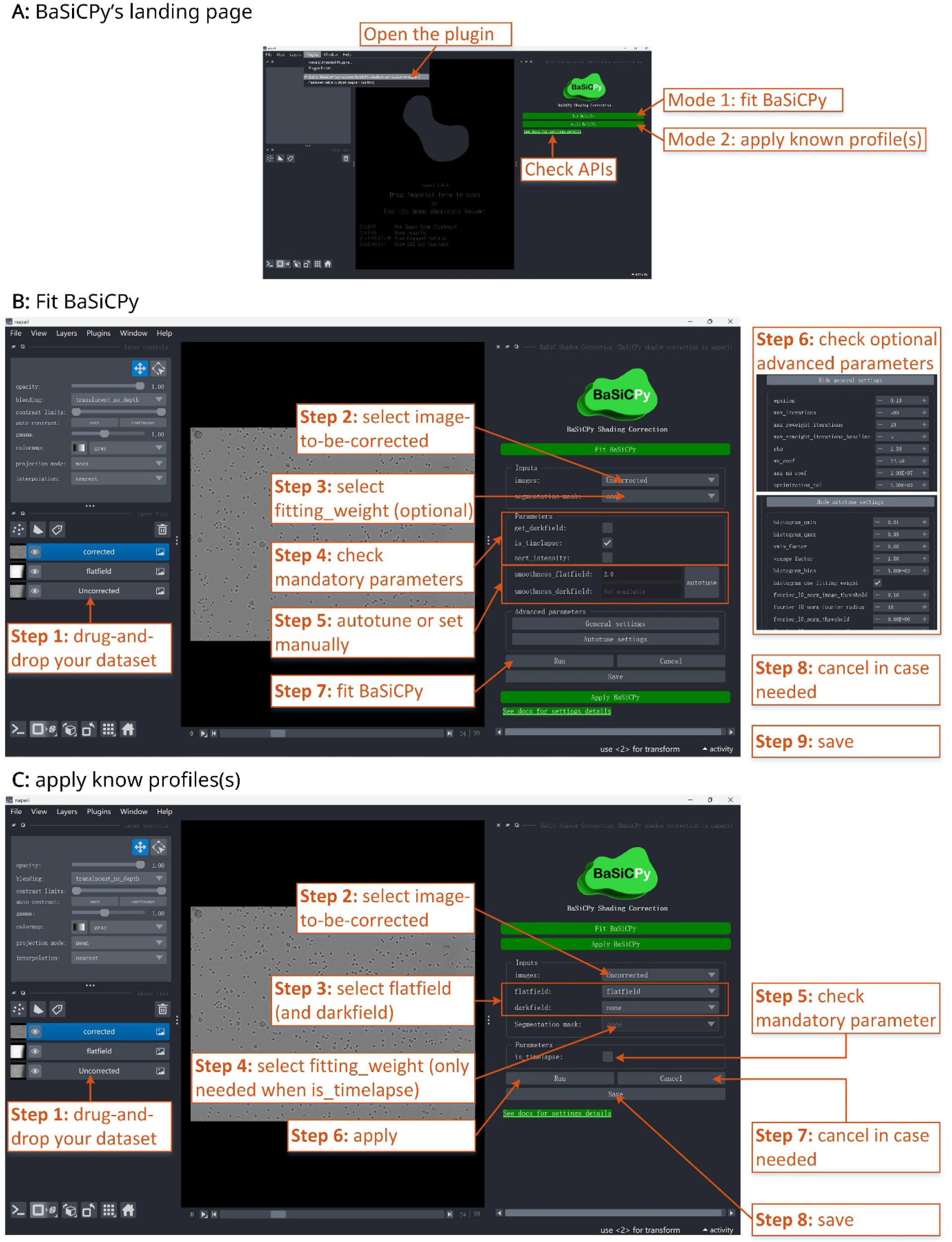
Graphical user interface (GUI) of the BaSiCPy napari plugin. (A) The BaSiCPy napari plugin offers two modes: Fit BaSiCPy (estimate illumination profiles) and Apply BaSiCPy (apply known profiles). (B) Workflow of “Fit BaSiCPy”. (C) Step-by-step workflow of “Apply BaSiCPy”.

**Extended Data Fig. 2.**
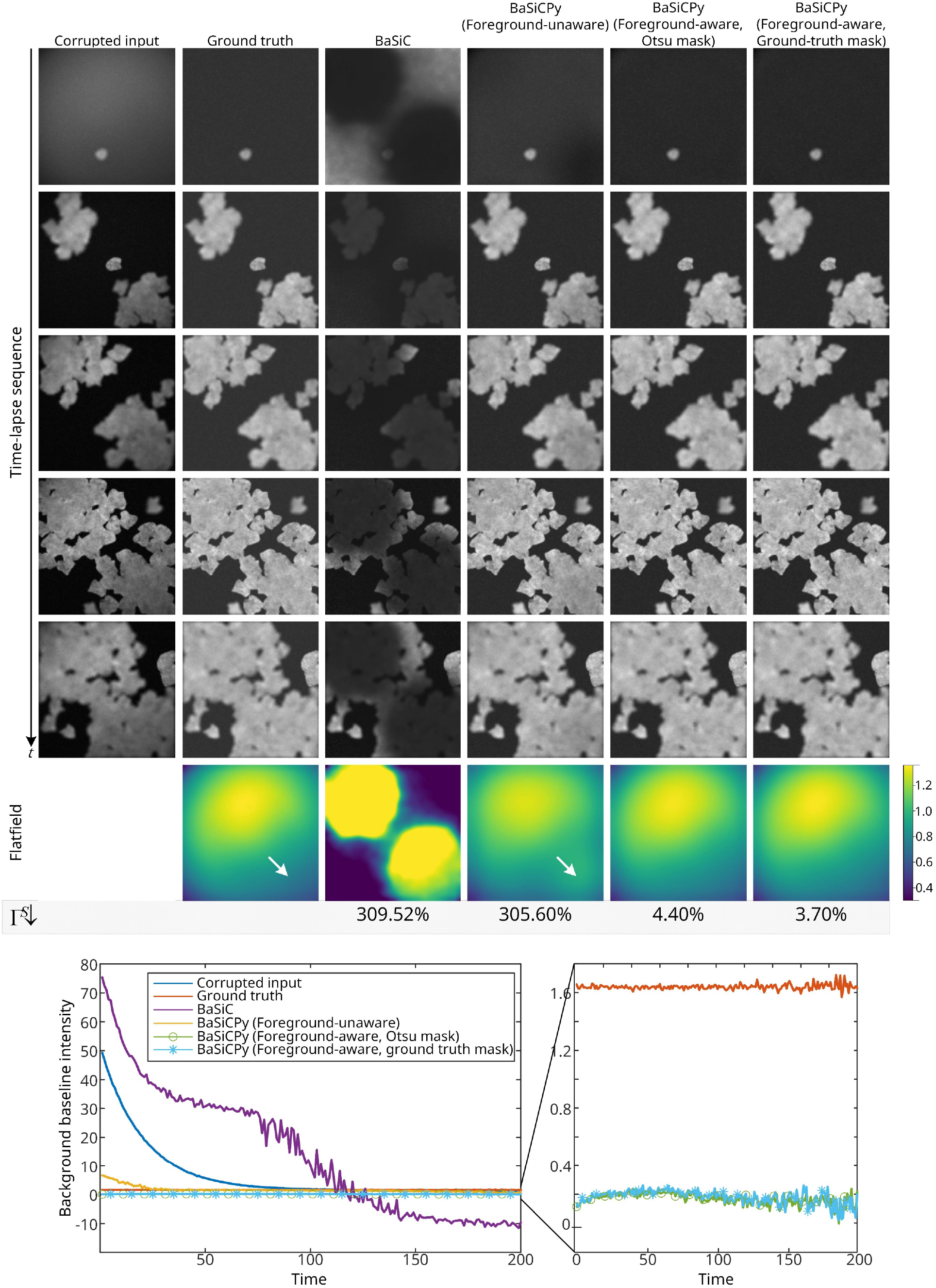
BaSiCPy successfully corrects spatially uneven illumination and temporal baseline drift in a cytoplasm dataset with correlated foreground. With BaSiCPy being foreground-aware, it accurately estimates the illumination profile without being biased by the correlated foreground. Moreover, BaSiCPy is particularly robust to the segmented foreground mask, yielding highly similar flatfields whether using the ground-truth foreground mask or a simple mask generated by Otsu thresholding. This is confirmed by Γ^*S*^, a measurement to quantify the similarity between the estimated and the ground-truth flatfields. In addition, BaSiCPy corrected the simulated exponential baseline drift over time. Note that foreground-unaware BaSiCPy (without mask guidance) outperforms original BaSiC due to its automatic hyperparameter selection (Autotune), yielding a lower Γ^*S*^. However, it remains susceptible to correlated foreground structures, as evidenced by residual hotspots (white arrows) in the estimated flatfield and a higher Γ^*S*^ compared to mask-guided estimates.

**Extended Data Fig. 3.**
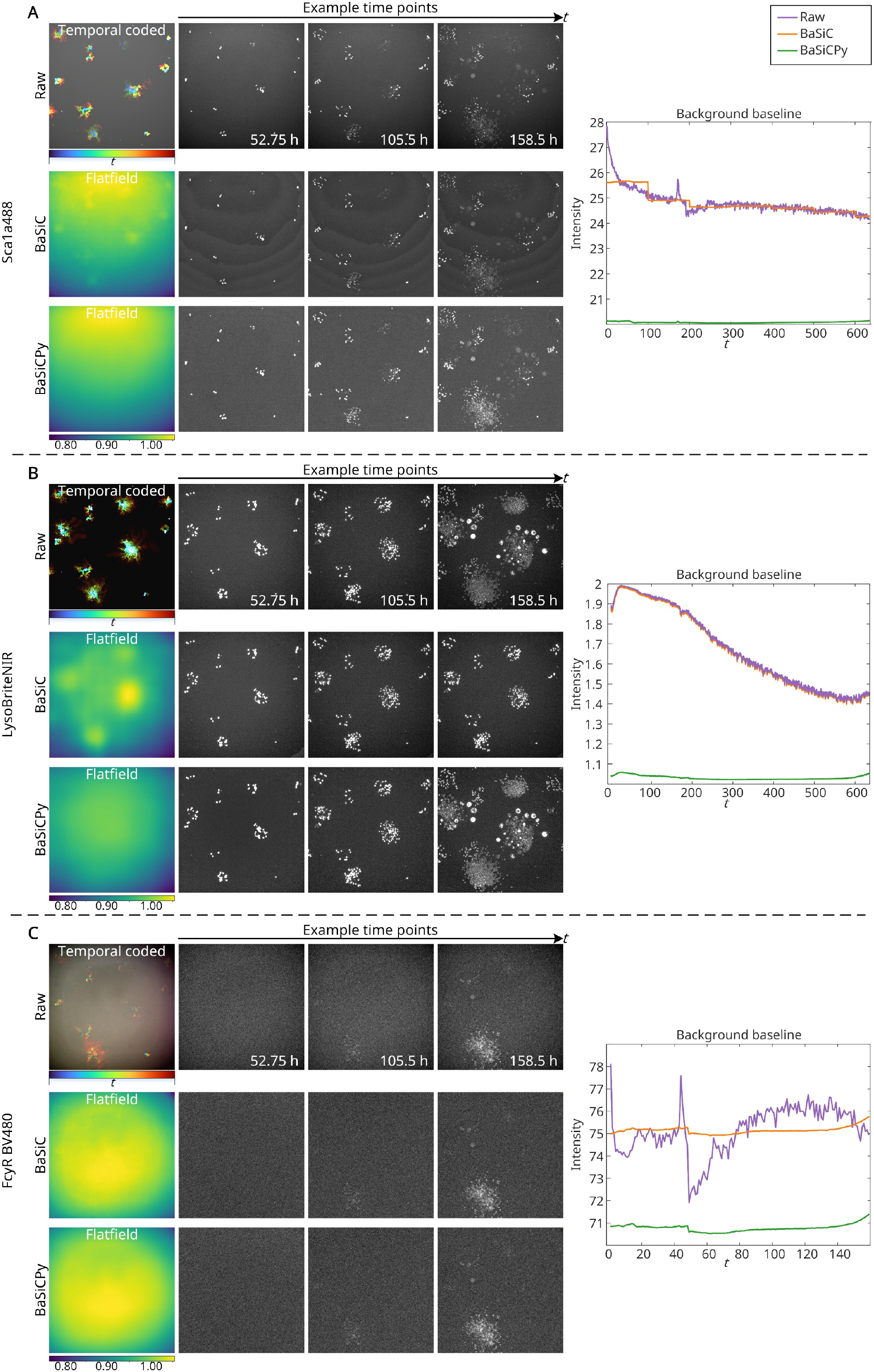
BaSiCPy corrects illumination inhomogeneity and temporal drift in real microscopy datasets across multiple channels. Representative results on three experimental channels (Sca1a488, LysoBriteNIR, and FcyR BV480) from the male mouse bone marrow hematopoietic stem cell dataset are shown. The result of the CD41 PE channel is shown in Fig. 2 of the main text. For each channel, the temporal-coded maximum intensity projection visualizes the overall time-lapse dynamics, followed by example frames at selected time points. BaSiCPy effectively removed spatial shading. The estimated flatfields (bottom panels) reveal that BaSiCPy accurately captures the underlying illumination profiles, while BaSiC (middle panels) shows residual bias or overfitting due to correlated structures. The rightmost plots quantify the background baseline over time, demonstrating that BaSiCPy yields temporally stable fluorescence intensity across the sequence and eliminates long-term intensity fluctuations across all tested channels.

**Extended Data Fig. 4.**
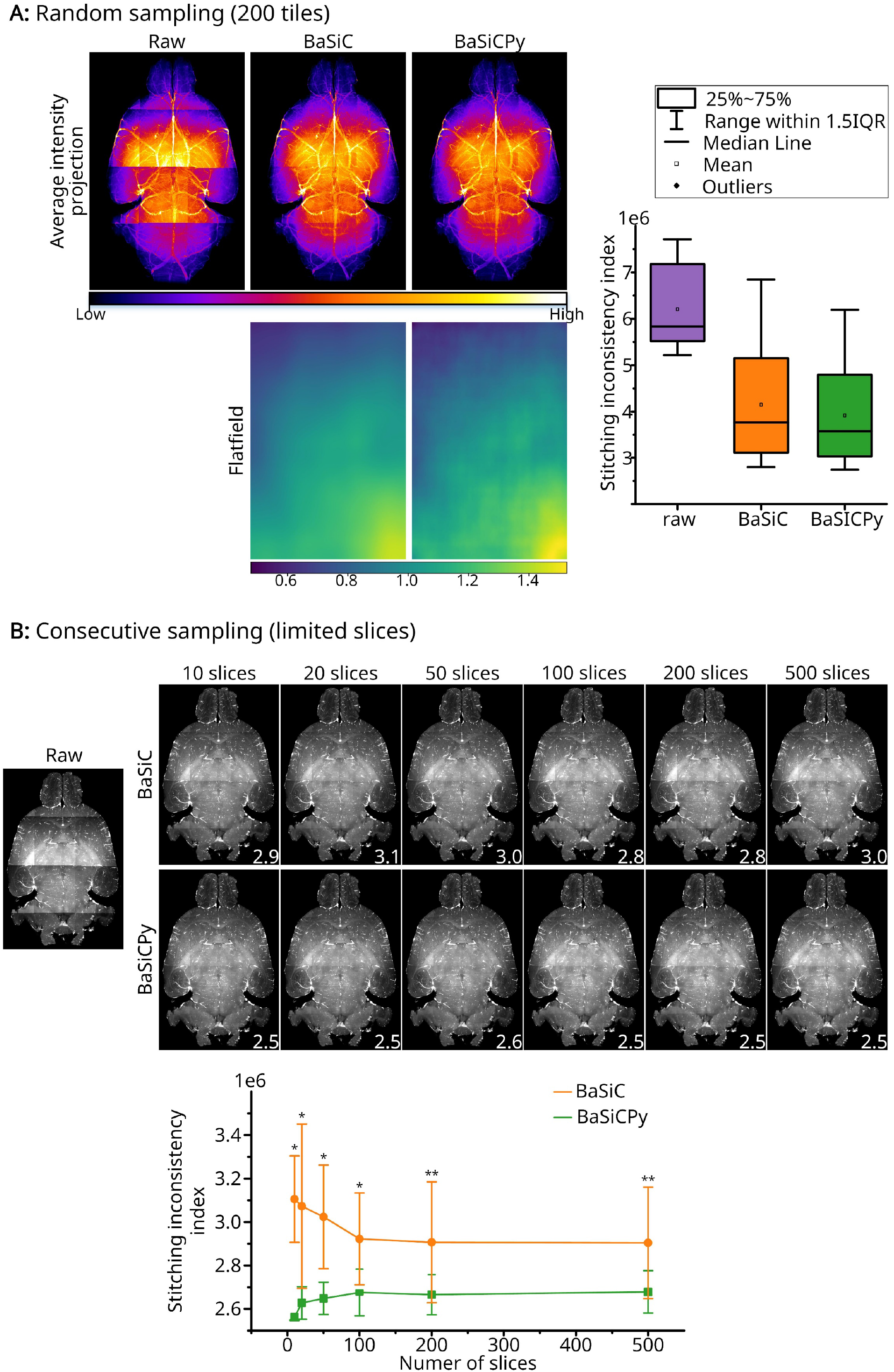
Sampling robustness of BaSiCPy under different flatfield estimation strategies. (A) Random sampling (200 tiles). Flatfield was estimated using 200 tiles randomly sampled across the entire mouse brain volume acquired with an UltraMicroscope II system. Under this well-sampled condition, BaSiC and BaSiCPy produced visually comparable corrections in the average intensity projection. The corresponding estimated flatfields are shown below. The boxplot shows lower stitching inconsistency values for BaSiCPy than for BaSiC under this sampling condition. (B) Consecutive sampling (limited slices). To mimic spatially constrained sampling, flatfield estimation was performed using only spatially consecutive slices (10–500 slices). While BaSiC exhibited unstable behaviors under limited sampling, BaSiCPy remained robust even with a small number of slices. (Line plot) Quantification of stitching inconsistency index as a function of the number of consecutive slices used for flatfield estimation. Error bars represent standard deviation. Significant differences between BaSiC and BaSiCPy at each sampling level are observed and indicated in the plot (*P < 0.05, **P < 0.01; paired Wilcoxon signed-rank test).

**Extended Data Fig. 5.**
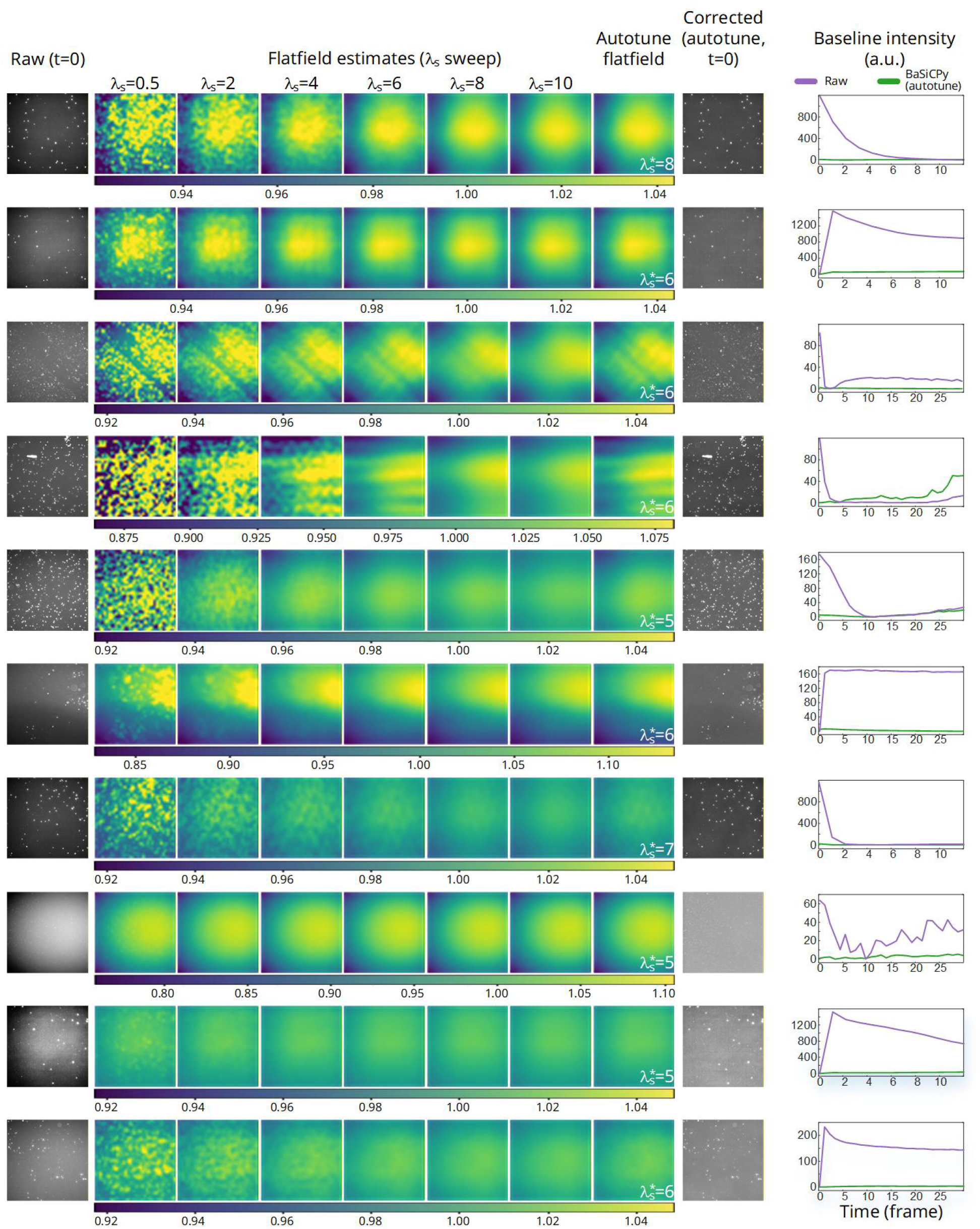
Autotune enables adaptive flatfield smoothness selection across heterogeneous wells in an antimicrobial single-cell testing (ASCT) time-lapse assay. Each row corresponds to one independent well acquired at different imaging positions under drug treatment conditions. For each well, flatfield estimation was performed using a range of smoothness parameters (*λ*_*s*_ = 0.5–10), alongside the Autotune-selected optimal parameter 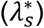. The selected 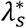 varies across wells, reflecting differences in foreground density, background levels, and illumination characteristics. Low *λ*_*s*_ (under-smoothing) introduces high-frequency artifacts into the estimated flatfield due to foreground leakage, whereas high *λ*_*s*_ (over-smoothing) fails to capture genuine illumination inhomogeneity. In contrast, Autotune selects well-adapted parameters that yield stable and visually consistent flatfield estimates without manual tuning. Right panels show the baseline intensity over time for PI fluorescence, demonstrating effective stabilization of temporal baseline drift across wells. Within each well, flatfield visualizations across different *λ*_*s*_ values share a common color scale. Y-axis scales of the baseline plots differ between wells due to varying baseline intensity ranges.

**Extended Data Fig. 6.**
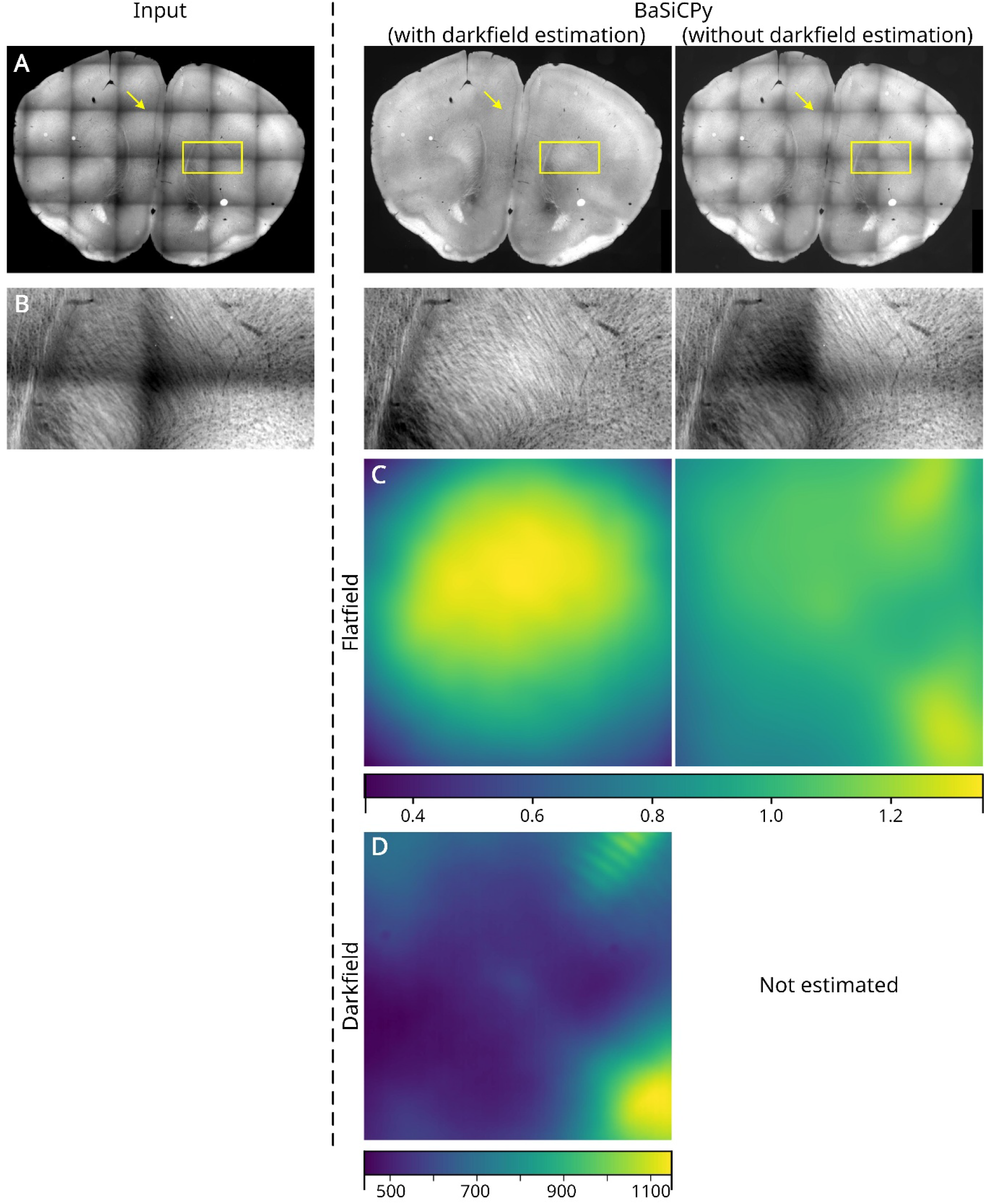
Impact of darkfield estimation in BaSiCPy. In general, darkfield is unnecessary in BaSiCPy and is disabled by default. In some edge cases, for example, this whole-slice imaging of a mouse brain, darkfield estimation can compensate for additive artifacts from stray light and residual excitation leakage through imperfect filters. (A) Only with darkfield estimation can the visible seams in the original image be removed (yellow arrows). A zoomed-in region is shown in (B). The estimated flatfield and darkfield (if applicable) are shown in (C) and (D), respectively. A more detailed discussion regarding this dataset can be found in the original BaSiC paper. Again, most modern acquisitions show a negligible additive background, so darkfield estimation is not recommended in general.

**Extended Data Fig. 7.**
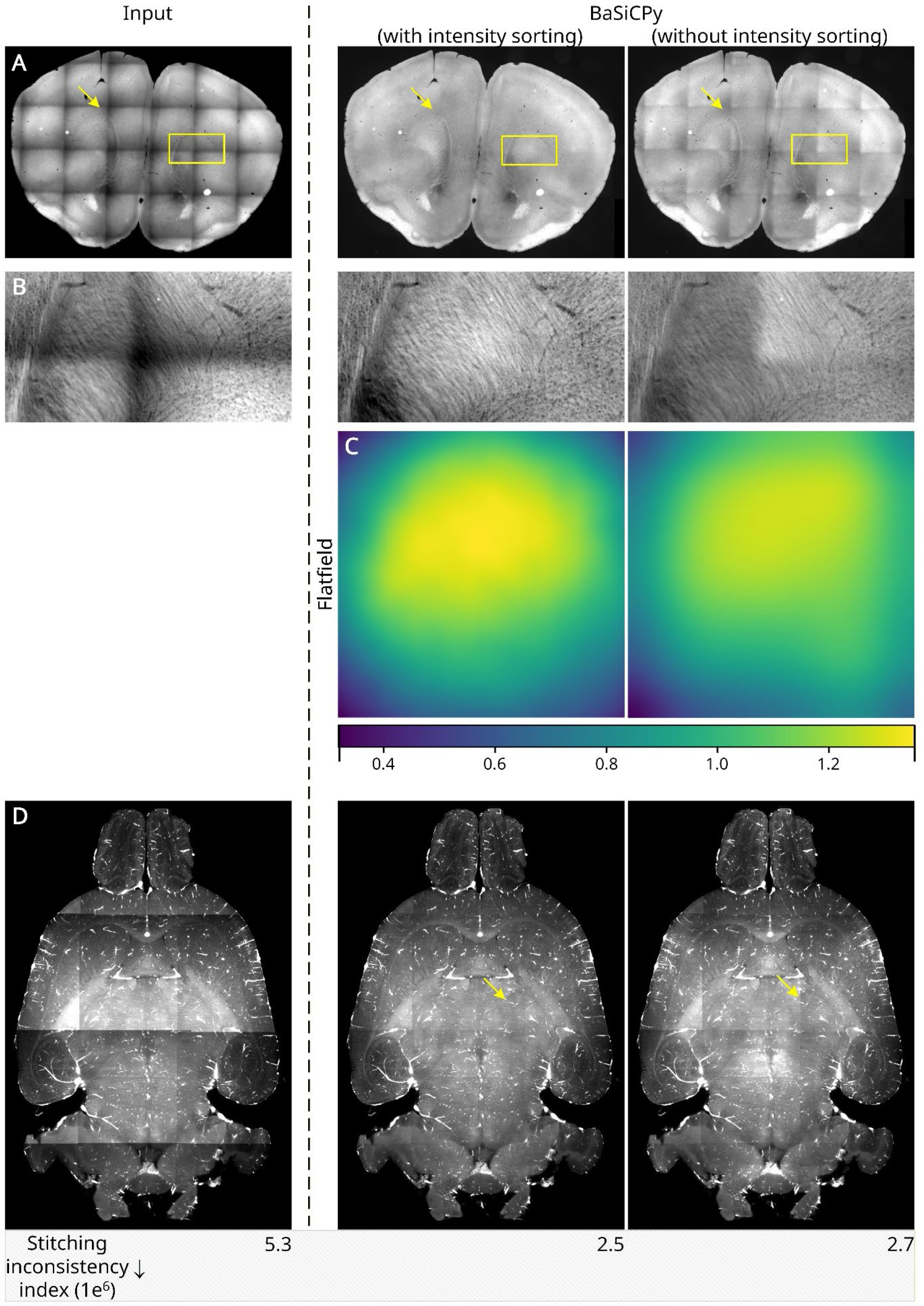
Impact of intensity sorting in BaSiCPy. (A) Input whole-slice imaging (WSI) dataset exhibiting pronounced tile-border intensity discontinuities. (B) Zoomed-in view of the region indicated by yellow boxes in (A). (C) Estimated flatfields obtained using BaSiCPy with intensity sorting (left) and without intensity sorting (right). Intensity sorting produces a smoother illumination profile and improved correction in this tissue dataset (see residual seams indicated by yellow arrows in (A)). (D) Results on a mouse brain dataset acquired using an UltraMicroscope II system (also shown in Extended Data Fig. 4). BaSiCPy with intensity sorting yields a lower residual stitching inconsistency (2.5 × 10^6^) compared to correction without sorting (2.7 × 10^6^). Yellow arrows indicate residual stitching artifacts observed after BaSiC correction but reduced by BaSiCPy.

**Extended Data Fig. 8.**
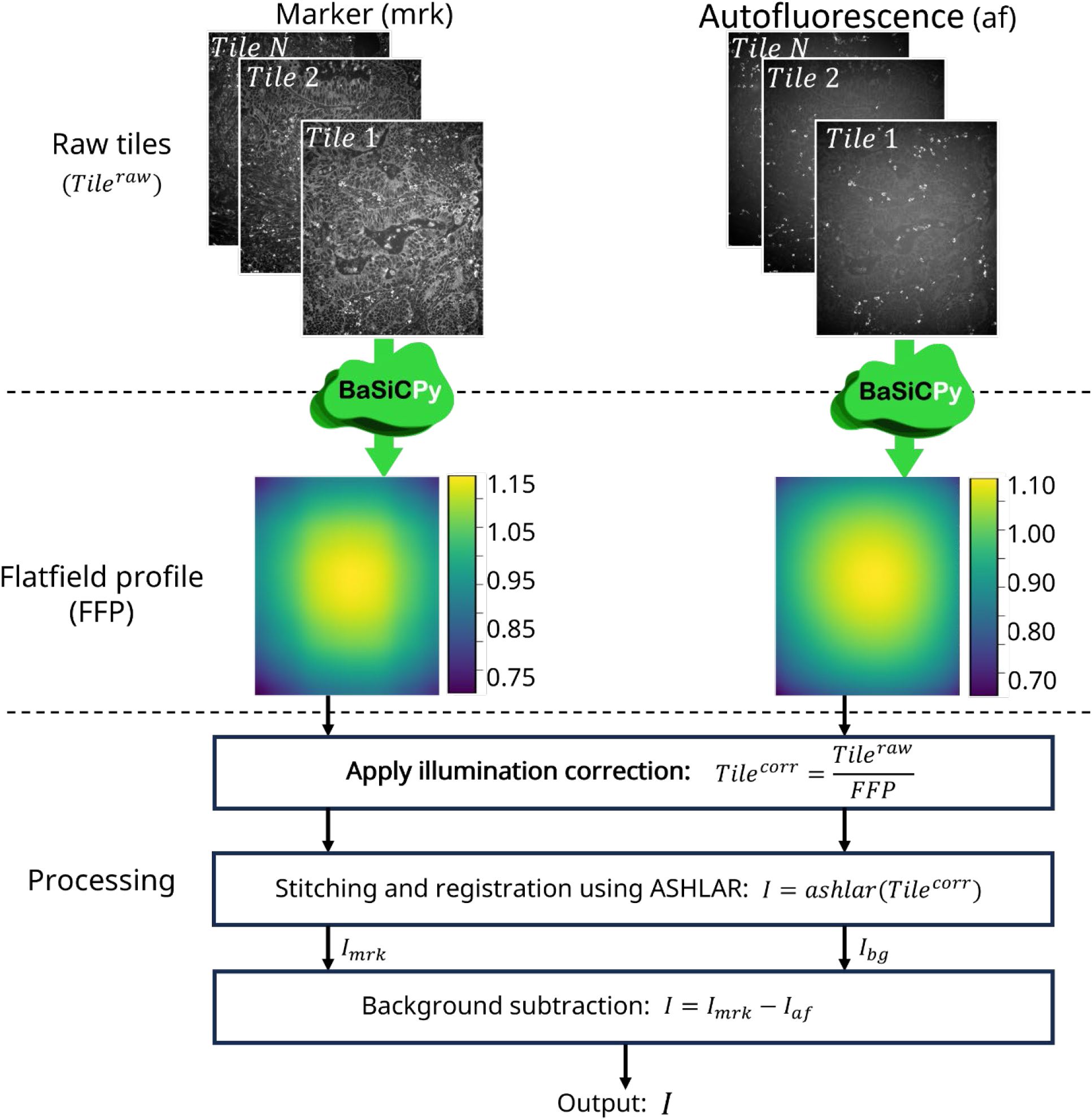
Flatfield estimation strategy for cyclic multiplexed imaging. In cyclic multiplexed workflows, marker and autofluorescence reference images are typically acquired in separate channels across multiple tiles and cycles. Owing to the spatially heterogeneous and cycle-dependent nature of autofluorescence, the background signal does not conform to a quasi-homogeneous darkfield component assumed by classical illumination correction models. We therefore recommend independent flatfield estimation and correction for marker tiles and autofluorescence reference tiles using BaSiCPy, followed by weighted autofluorescence subtraction at the stitched image level. This channel-wise correction preserves the validity of the multiplicative flatfield model and avoids bias introduced by directly applying illumination correction to marker–autofluorescence difference images.

**Extended Data Fig. 9.**
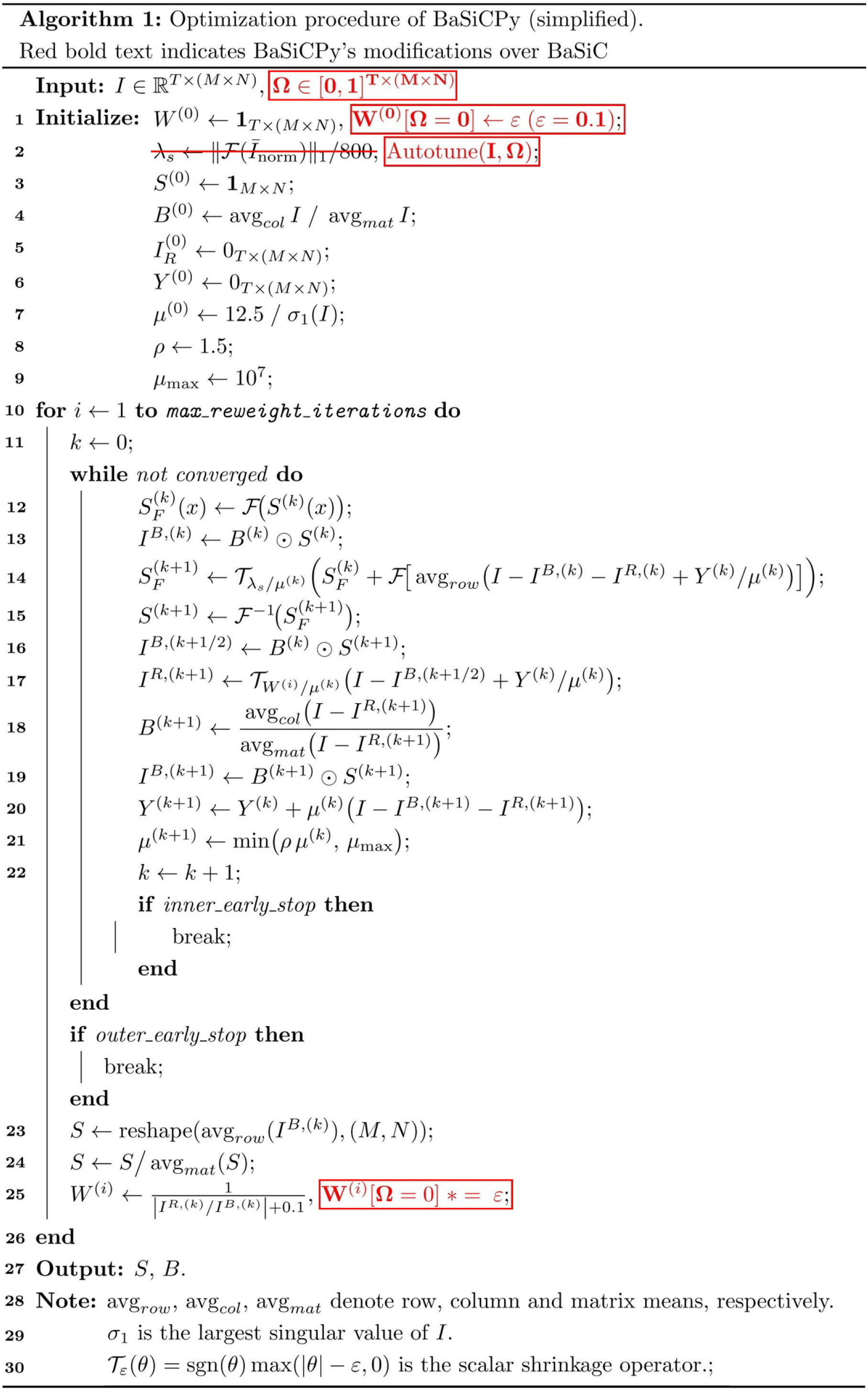
Schematic overview of BaSiCPy modifications to BaSiC. The pseudocode illustrates where the two BaSiCPy modifications are introduced within the BaSiC workflow. The original BaSiC operations are shown in black, whereas the BaSiCPy-specific modifications are highlighted in red bold text. First, a foreground-aware segmentation mask Ω is introduced to occlude the influence of correlated foreground on illumination profile estimation. Second, balancing parameter *λ*_*s*_, which is critical in BaSiCPy in terms of performance, is automatically tuned using Autotune operator. Note that the pseudocode provides an approximate overview of the iterative optimization; certain implementation-specific operations, such as estimating illumination profiles in a downsampled space to accelerate computation, are inherited from BaSiC^2^ and omitted here for clarity.

**Extended Data Fig. 10.**
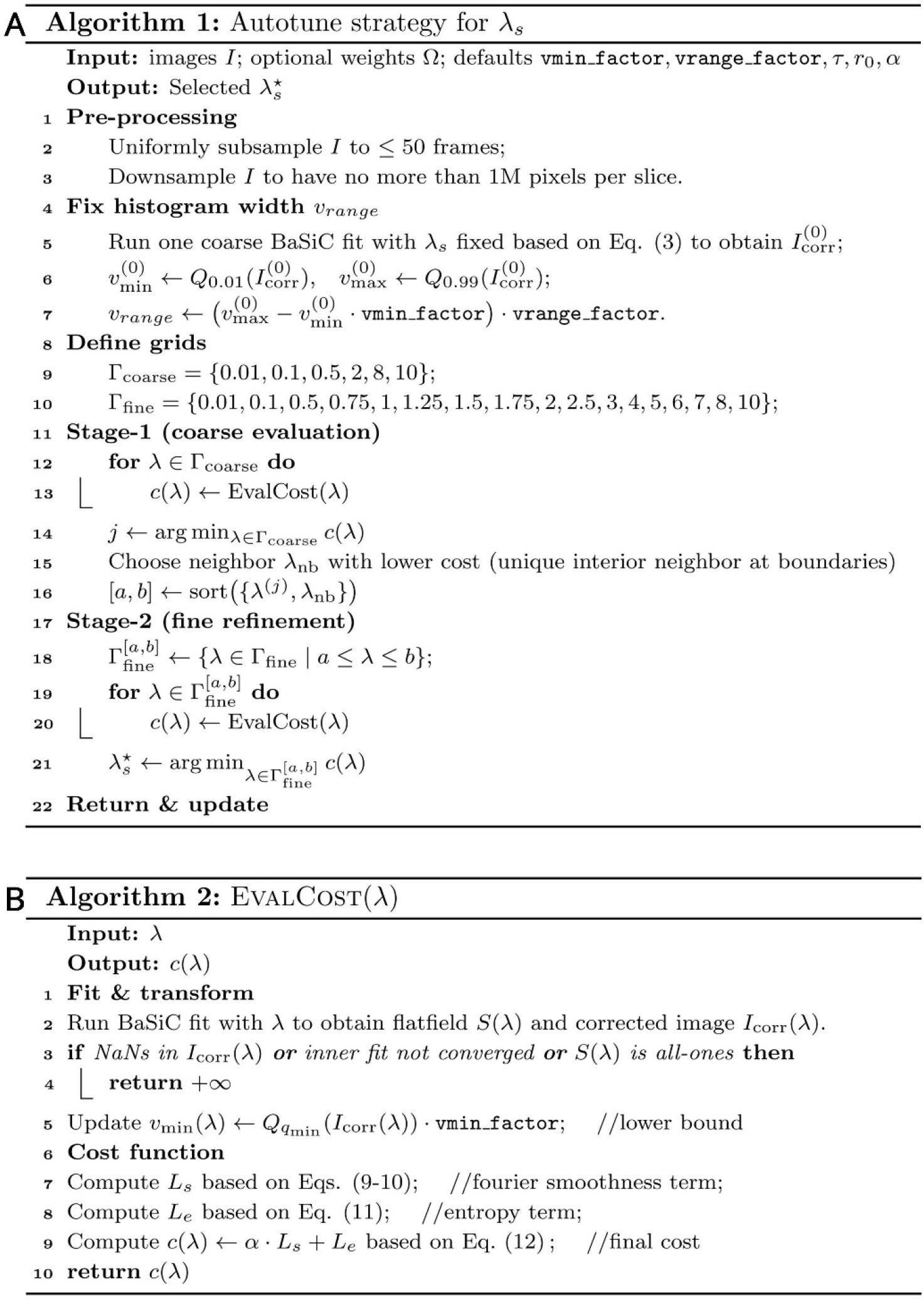
Autotune in BaSiCPy. (A) main workflow of Autotune strategy in BaSiCPy using a two-stage grid search to select the best hyperparameter *λ*_*s*_. For each candidate *λ*, a cost function that balances between the Fourier-based smoothness term and the entropy-based compactness term is detailed in (B).

## Extended Data Tables

**Extended Data Table 1.**
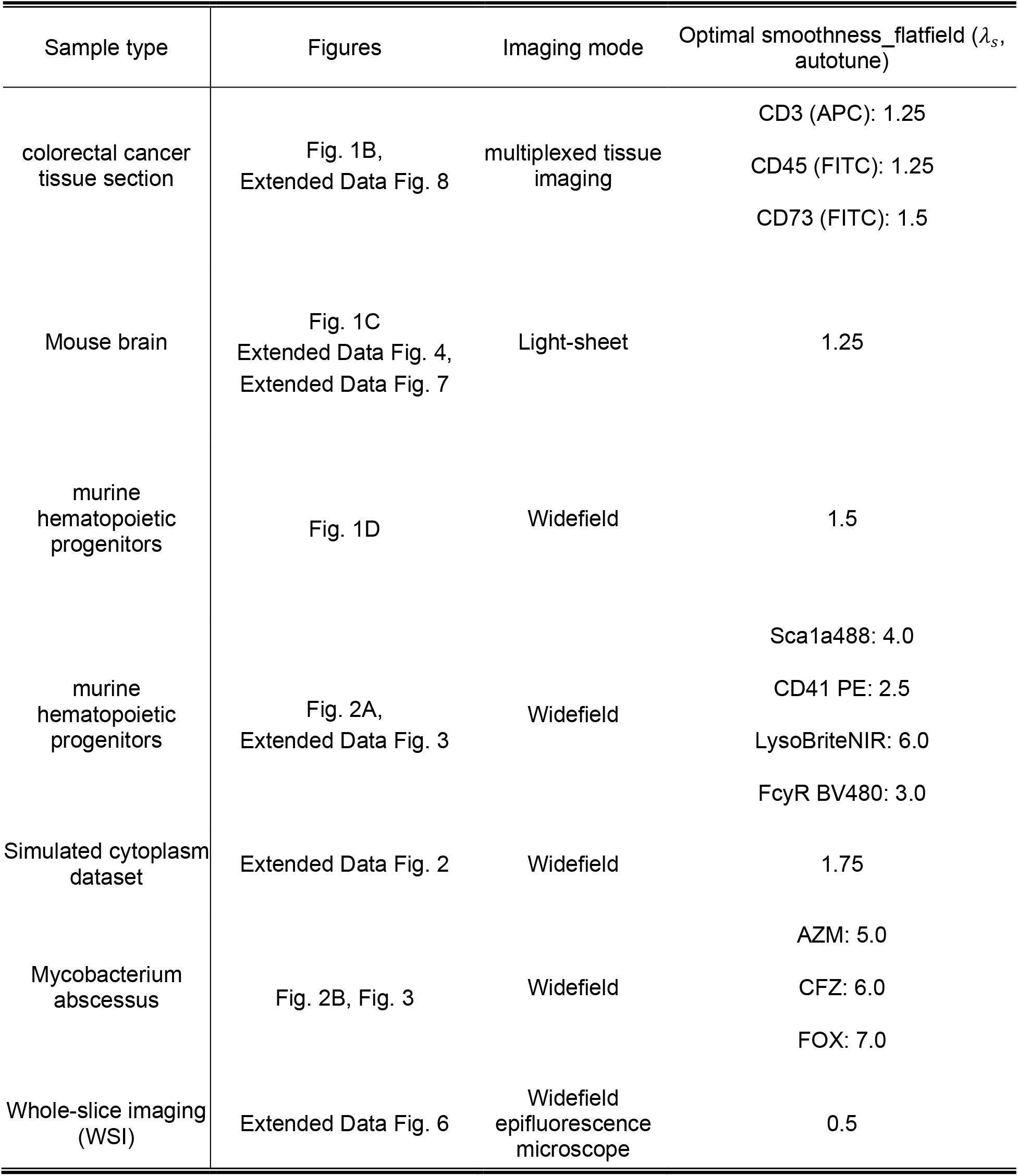
Dataset-dependent variability of optimal flatfield smoothness necessitates automated parameter selection.

| Sample type | Figures | Imaging mode | Optimal smoothness_flatfield ($\lambda_s$, autotune) |
| --- | --- | --- | --- |
| colorectal cancer tissue section | Fig. 1B,<br>Extended Data Fig. 8 | multiplexed tissue imaging | CD3 (APC): 1.25<br>CD45 (FITC): 1.25<br>CD73 (FITC): 1.5 |
| Mouse brain | Fig. 1C<br>Extended Data Fig. 4,<br>Extended Data Fig. 7 | Light-sheet | 1.25 |
| murine hematopoietic progenitors | Fig. 1D | Widefield | 1.5 |
| murine hematopoietic progenitors | Fig. 2A,<br>Extended Data Fig. 3 | Widefield | Sca1a488: 4.0<br>CD41 PE: 2.5<br>LysoBriteNIR: 6.0<br>FcyR BV480: 3.0 |
| Simulated cytoplasm dataset | Extended Data Fig. 2 | Widefield | 1.75 |
| Mycobacterium abscessus | Fig. 2B, Fig. 3 | Widefield | AZM: 5.0<br>CFZ: 6.0<br>FOX: 7.0 |
| Whole-slice imaging (WSI) | Extended Data Fig. 6 | Widefield epifluorescence microscope | 0.5 |

**Extended Data Table 2.** Manual hyper-parameters used by different datasets.

| Sample type | Figures | Is timelapse | Intensity sorting | Foreground-informed | Darkfield enabled |
| --- | --- | --- | --- | --- | --- |
| colorectal cancer tissue section | Fig. 1B, Extended Data Fig. 8 | X | X/✓ | X | X |
| Mouse brain | Fig. 1C<br>Extended Data Fig. 4,<br>Extended Data Fig. 7 | X | ✓ | ✓ | X |
| murine hematopoietic progenitors | Fig. 1D | ✓ | X | ✓ | X |
| murine hematopoietic progenitors | Fig. 2A,<br>Extended Data Fig. 3 | ✓ | X | ✓ | X |
| Mycobacterium abscessus | Fig. 2B, Fig. 3 | ✓ | X | X | X |
| Simulated cytoplasm dataset | Extended Data Fig. 2 | ✓ | X | ✓ | X |
| Whole-slice imaging (WSI) | Extended Data Fig. 6 | X | ✓ | X | ✓ |

## Notes

https://basicpy.readthedocs.io/en/latest/

https://github.com/peng-lab/BaSiCPy

https://zenodo.org/records/6334810

### Summary of Updates

The initial version (v1) was prepared and submitted under the responsibility of Yu Liu and Tingying Peng only. This revised version includes corrections and updates to the author information, as well as improvements to the manuscript's clarity and presentation.

## References

1. Schapiro, D. et al. MCMICRO: a scalable, modular image-processing pipeline for multiplexed tissue imaging. Nat. Methods 19, 311–315 (2022).

2. Peng, T. et al. A BaSiC tool for background and shading correction of optical microscopy images. Nat. Commun. 8, 14836 (2017).

3. Smith, K. et al. CIDRE: an illumination-correction method for optical microscopy. Nat. Methods 12, 404–406 (2015).

4. Singh, S., Bray, M. -A., Jones, T. R. & Carpenter, A. E. Pipeline for illumination correction of images for high-throughput microscopy. J. Microsc. 256, 231–236 (2014).

5. Pozniak, J. et al. A TCF4-dependent gene regulatory network confers resistance to immunotherapy in melanoma. Cell 187, 166–183.e25 (2024).

6. Gupta, D. et al. Panoramic visual statistics shape retina-wide organization of receptive fields. Nat. Neurosci. 26, 606–614 (2023).

7. Thornton, M. A. et al. Long-term in vivo three-photon imaging reveals region-specific differences in healthy and regenerative oligodendrogenesis. Nat. Neurosci. 27, 846–861 (2024).

8. Jovanovic, A. et al. Large-scale testing of antimicrobial lethality at single-cell resolution predicts mycobacterial infection outcomes. Nat. Microbiol. 11, 566–583 (2026).

9. Lehmussola, A., Ruusuvuori, P., Selinummi, J., Huttunen, H. & Yli-Harja, O. Computational Framework for Simulating Fluorescence Microscope Images With Cell Populations. IEEE Trans. Med. Imaging 26, 1010–1016 (2007).

10. Lin, J., Fallahi-Sichani, M., Chen, J. & Sorger, P. K. Cyclic Immunofluorescence (CycIF), A Highly Multiplexed Method for Single-cell Imaging. Curr. Protoc. Chem. Biol. 8, 251–264 (2016).

11. Kinkhabwala, A. et al. MACSima imaging cyclic staining (MICS) technology reveals combinatorial target pairs for CAR T cell treatment of solid tumors. Sci. Rep. 12, 1911 (2022).

12. Black, S. et al. CODEX multiplexed tissue imaging with DNA-conjugated antibodies. Nat. Protoc. 16, 3802–3835 (2021).

13. Muhlich, J. L. et al. Stitching and registering highly multiplexed whole-slide images of tissues and tumors using ASHLAR. Bioinformatics 38, 4613–4621 (2022).

